# The Australian Genetics of Depression Study: Study Description and Sample Characteristics

**DOI:** 10.1101/626762

**Authors:** Enda M Byrne, Katherine M Kirk, Sarah E Medland, John J McGrath, Richard Parker, Simone Cross, Lenore Sullivan, Dixie J Statham, Douglas F Levinson, Julio Licinio, Naomi R Wray, Ian B Hickie, Nicholas G Martin

**Affiliations:** Institute for Molecular Bioscience, University of Queensland, Brisbane, Australia; QIMR Berghofer Institute of Medical Research, Brisbane, Australia; Queensland Brain Institute, University of Queensland, Brisbane, Australia; Queensland Centre for Mental Health Research, The Park Centre for Mental Health, Wacol, Australia; National Center for Register-based Research, University of Aarhus, Aarhus, Denmark; Federation University, Ballarat, Australia; Department of Psychiatry, Stanford University School of Medicine, Palo Alto, CA, USA; College of Medicine, State University of New York Upstate Medical University, Syracuse, NY, USA; Department of Psychiatry, State University of New York Upstate Medical University, Syracuse, NY, USA; Brain and Mind Centre, The University of Sydney, Sydney, Australia

**Keywords:** depression, genetics, GWAS, antidepressants

## Abstract

**Objectives:** Depression is the most common psychiatric disorder and the largest contributor to global disability. The Australian Genetics of Depression study was established to recruit a large cohort of individuals who have been diagnosed with depression, and to investigate genetic and environmental risk factors for depression and response to commonly prescribed antidepressants. This paper describes the recruitment and characteristics of the sample.

**Methods:** Participants completed an online questionnaire that consisted of a compulsory module that assessed self-reported psychiatric history, clinical depression using the Composite Interview Diagnostic Interview Short Form, and experiences of using commonly prescribed antidepressants. Further voluntary modules assessed a wide range of traits of relevance to psychopathology. Participants who reported they were willing to provide a DNA sample were sent a saliva kit in the mail.

**Results:** A total of 20,689 participants, 75% of whom were female, enrolled in the study. The average age of participants was 43 years ± 15 years. 15,807 participants (76% of the participant group) returned saliva kits. The overwhelming majority of participants reported being given a diagnosis of depression by a medical practitioner and 88% met the criteria for a depressive episode. Rates of comorbidity with other psychiatric disorders were high. Two-thirds of the sample reported having taken more than one type of antidepressant during treatment for their depression.

**Conclusions:** This study was effective in recruiting a large community sample of people with a history of clinical depression, highlighting the willingness of Australians to engage with medical research. A combination of recruitment through health records and media as well as use of an online questionnaire made it feasible to recruit the large sample needed for investigating the genetics of common diseases. It will be a valuable resource for investigating risk factors for depression, treatment response to antidepressants and susceptibility to side effects.

## Introduction

Approximately 20% of Australians will be diagnosed with a depressive disorder in their lifetime. As a consequence of this high prevalence, impact on function and risk to later ill-health and premature death, depressive disorders contribute the largest burden of disease due to common mental disorders (Whiteford et al., 2013; Ferrari et al., 2013) and place a substantial burden on the economy in terms of days lost to disability.

Among psychiatric disorders, depression is moderately heritable, with approximately 40% of the variance in liability to depression attributable to genetic factors (Sullivan et al., 2000). Initial efforts to identify depression risk variants using genome-wide association studies (GWAS) did not bear fruit due to insufficient power (Wray et al., 2012). Common genetic variants for psychiatric disorders have small effect sizes and hence robust detection requires sample sizes in the tens of thousands of individuals in order to robustly to detect them. Substantial progress has been made in the last few years in identifying genetic variants that increase risk to depressive symptoms and major depression (Wray et al., 2018; Howard et al., 2018; consortium, 2015). These discoveries have been facilitated by the collaboration of researchers worldwide in the Psychiatric Genomics Consortium (PGC). The most recent GWAS for depression which included data from the PGC, the personal genetics company 23andMe, the UK Biobank, and DeCODE, identified 102 independent genetic variants that increase risk of depression (Howard et al., 2019). The identified variants explain only a fraction of the overall liability and larger studies are needed to identify more individual variants and to improve the predictive power of polygenic risk scores. Thus, the psychiatric genomics community aims to collect data on 1 million cases with depression in order to elucidate the genetics.

Antidepressants are a frontline treatment for moderate to severe depression (Malhi and Mann, 2018), but do not provide benefit for all patients and have side effects, leading to poor adherence and reduced quality of life. Variability in response to antidepressants and experiencing side effects have a poorly understood genetic component (Tansey et al., 2013; Hodgson et al., 2014). As they are one of the most commonly prescribed medications and many individuals are exposed to several different drugs, or drug classes, before symptoms improve, there is an urgent need to understand the reasons for such wide individual variability in therapeutic response and the experience of side effects. Results from pharmacogenetic studies of response and side effects have been mixed, likely because of insufficient sample sizes (Biernacka et al., 2016; Uher et al., 2010; Investigators et al., 2013; Tansey et al., 2012; Li et al., 2016).

To identify genetic and non-genetic risk factors for depression risk, antidepressant response and side-effects, we established the Australian Genetics of Depression Study (AGDS). By approaching those using antidepressants through the Pharmaceutical Benefits Scheme and those who have been treated for depression through a media campaign, we aimed to recruit 10,000 cases with depression to an online study and obtain a DNA sample using a saliva kit to contribute to the wider PGC effort to identify genetic variants that increase risk to depression and antidepressant response. Here we describe the aims of the study, the genetic and phenotype data collection procedures and the characteristics of the sample.

## METHODS

### Participant Recruitment

Participants were recruited to the Australian Genetics of Depression Study (www.geneticsofdepression.org.au) using two separate approaches: (i) recruitment based on nationwide, pharmaceutical prescription history in the last 4.5 years and (ii) a media publicity campaign. A schematic of the design and aims of the study is shown in Figure 1.

**Figure 1.**
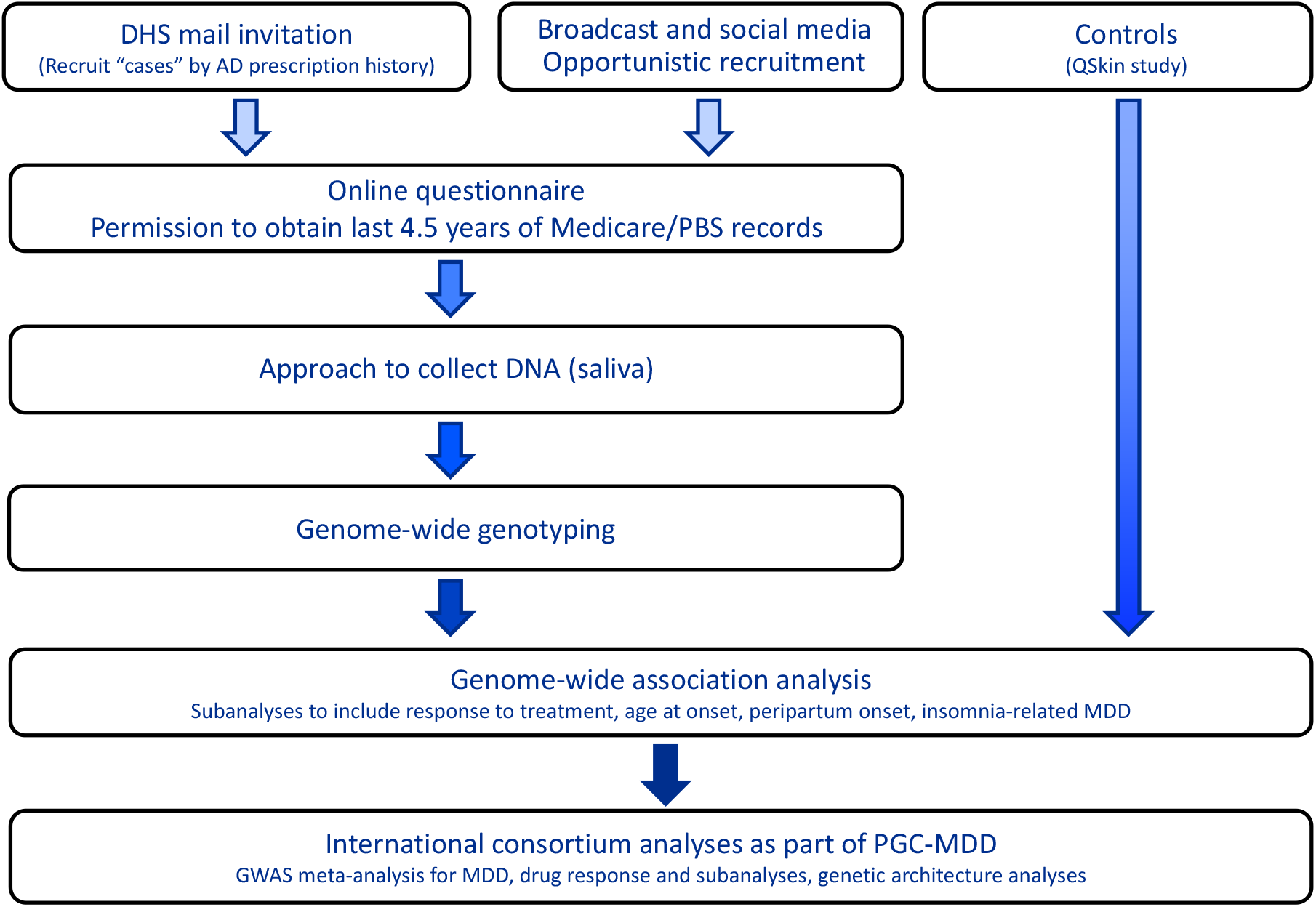
Schematic of the Australian Genetics of Depression Study

#### Recruitment via pharmaceutical prescription history

The Australian Government subsidises certain healthcare services through the Medicare Benefits Scheme (MBS) and prescription medications through the Pharmaceutical Benefits Scheme (PBS). Records for the most recent 4.5 years’ services provided are retained by the Australian Government Department of Human Services (DHS). While these records are not accessible to researchers for the purposes of identifying potential research study participants, DHS is able to send invitations on behalf of researchers to individuals meeting specific selection criteria.

After receiving approval from the DHS research ethics committee, two waves of recruitment were undertaken using this method. A pilot study in which DHS sent 10,000 invitation letters to Australian residents aged 18-30 who had received four or more prescriptions in the previous 4.5 years for any of the 10 most commonly prescribed antidepressant medications (single medication or a combination) was initiated in September 2016. Only community patients were selected; individuals with residential locations in the PBS database corresponding to hospitals, aged-care facilities and correctional facilities were excluded. This group of invitees was 65% female, reflecting the higher prevalence of depression in women. Potential participants were sent a letter by the DHS explaining that were being contacted on behalf of researchers at QIMR Berghofer to participate in a study of the genetics of depression. The letter provided details of the study website and also a phone number that they could contact for more information. A total of 294 individuals responded to this invitation and enrolled in the study.

The second DHS-based recruitment wave started in April 2017 and involved sending 100,000 invitation letters to people selected using similar selection criteria to the pilot study, except that the upper age restriction for participants was removed.

#### Recruitment through Media Publicity Campaign

A Sydney-based PR company specialising in health sector campaigns (VIVA! Communications) was contracted to manage the media campaign, which was launched on April 4 2017 and utilised a combination of national broadcast, print and social media to promote knowledge of and interest in the study among the general community. This coincided with the second wave of recruitment through DHS. The campaign encouraged participation among “Australian adults who have, or are continuing to be treated for clinical depression by a doctor, psychologist or psychiatrist”. A second wave of the media campaign was initiated 6 months after the initial one in September 2017.

### Study Design

#### Enrolment

In both the DHS recruitment letter and the media public appeal, potential participants were asked to go to the study website which was hosted on the secure QIMR Berghofer server. Upon going to the website, the information sheet which provided details of the aims of the study as well as a consent form were available for viewing. The information sheet provided telephone and e-mail contact details for the study co-ordinator and institute ethics committee in case participants had any questions. Those not interested in participating were provided with simple instructions on how to exit. The identity of potential participants was not known to the researchers prior to their decision to enrol in the study. The DHS did not provide identifying information to the research team on who was mailed. Before being asked to provide any identifying information, prospective participants were asked to confirm that they had read and understood the information sheet and also to confirm that they would be willing to provide a saliva sample.

Upon confirming that they would like to enrol in the study, participants were asked to provide their name, age and contact details which were stored securely on the QIMR server. After providing these details, each participant was assigned a unique link to the questionnaire which was hosted on the Qualtrics website. This transition between websites was seamless to the participant.

#### Access to Medicare and PBS records

Participants were also asked to consent to provide access to their list of Medicare and Pharmaceutical Benefits Scheme records for the previous 4.5 years, and approximately 75% of participants did so. This consent process was separate to the overall consent to participate in the study, and participants could still enrol in the study without allowing access to these records. The consent form had to conform to the requirements of the Department of Human Services. Participants were shown an example of what MBS and PBS records look like prior to consenting so they would know what information would be available to researchers. Within the MBS and PBS data, the identifiers for the providing doctor, medical service or pharmacy are randomised so the provider and location are protected. It is possible to identify repeated claims from the same provider but not who the provider is.

#### Ethics

All study protocols were approved by the QIMR Berghofer Medical Research Institute Human Research Ethics Committee. The protocol for approaching participants through the DHS, enrolling them in the study, and consenting for accessing MBS and PBS records was approved by the Ethics Department of the Department of Human Services.

#### Saliva collection and DNA extraction

Several brands of saliva DNA kits were tested for suitability for use, including cost, ease of handling, and yield and quality of extracted DNA. The Isohelix GeneFix™ GFX-02 2mL saliva collector was selected due it being the most compact, reliable, easy to use, lightweight and therefore the least expensive to mail to participants.

After completing the core module of the questionnaire, participants were emailed to confirm their delivery address and their readiness to receive a saliva DNA kit. Upon confirmation, they were mailed a spit kit, together with a consent form to be signed and returned with the tube. We found that this confirmation step markedly increased compliance. Saliva samples were returned by study participants by pre-paid post. If the kit was not returned after 2 months, study personnel followed up by phone or email in order to maximise return rates. Upon return of the kit, DNA was extracted from the saliva sample and stored in freezers.

Genotyping is being conducted using the Illumina Global Screening Array 2.0 (GSA) and is expected to be completed mid 2019. GSA was developed by human genetic disease researchers to maximise utility for gene-mapping. It includes a common variant backbone component that maximises information for imputation of common variants in multiple ethnic populations as well as a suite of common and rare variants selected for known or likely association with a range of genetic disorders. Importantly for the purposes of this study, it includes several genetic variants with known pharmacogenetic associations from the Pharmacogenomics Knowledgebase (PharmGKB).

#### Controls – the QSkin study

The primary aim of the AGDS was to recruit as many individuals with depression as possible. There was no publicity initiated to recruit controls because an appropriate control sample is available from the QSkin study. QSkin was established in 2010 to investigate risk factors for melanoma and other skin cancers in a randomly sampled cohort of individuals aged between (40-69 years) from the state of Queensland (Olsen et al., 2012). To date, more than 40,000 participants have enrolled in QSkin. Recently, a genetics arm of the study was initiated and follows a similar protocol for collection of DNA using saliva kits returned by mail. At the time of saliva collection, participants are asked about their medical history, including whether they have ever been diagnosed with or treated for depression, bipolar disorder, schizophrenia/psychosis, anxiety, obsessive compulsive disorder, bulimia, anorexia nervosa, autism or ADHD. In addition, women are asked if they experienced either antenatal or postnatal depression. Moreover, participants were consented for access to MBS and PBS records which will permit screening for use of antidepressants in addition to the disease checklist screening items above.

More than 18,000 participants have been genotyped on the same SNP microarray chip – the Illumina GSA - and the genotype data will be merged with the AGDS study prior to genome-wide imputation. The QSKIN study thus provides a large sample of Australian controls selected at random from the population and genotyped on the same SNP chip

#### Questionnaire

The content of the Australian Genetics of Depression Study online questionnaire was developed over a period of 19 months between January 2015 and September 2016. The object was to maximise the amount of clinically relevant information collected with the shortest time commitment required of participants. To this end, we utilised a modular structure (Figure 2), with a core module eliciting essential information on self-report mental health diagnoses, medication response and side effects, depression diagnosis using the relevant section from the Composite International Diagnostic Interview (CIDI), screens for suicidality, mania and psychosis, and a question about family history of psychiatric disorders. Several psychiatrists in Australia and internationally with expertise in gene mapping studies and in studies of antidepressant response were consulted about the content of the questionnaire.

**Figure 2.**
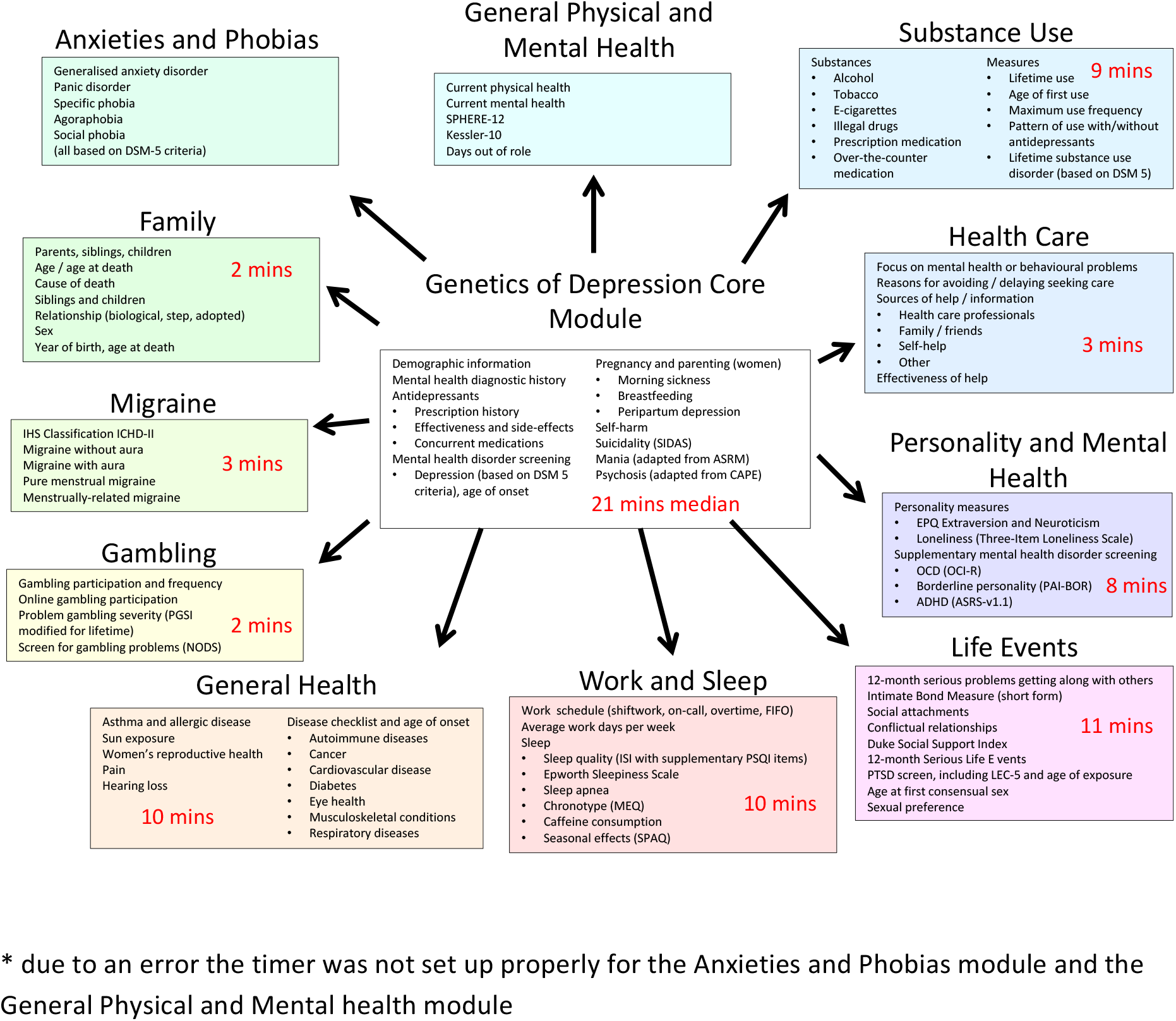
Overview of the structure and content of the AGDS questionnaire with median amount of time taken to complete each module*.

Ten additional “satellite” modules assessed a range of complex traits of relevance to mental health using a variety of scales and questionnaires (Figure 2). One module screened for clinical anxiety using the CIDI. The questionnaire was administered online using the Qualtrics™ software. Responses to individual questionnaire items were only required for items critical to phrasing of future questionnaire items and skip functionality (e.g. age, sex, number of children). The satellite modules could be completed in any order the participant chose once they had completed the core module. Participants were able to leave the survey and return at their convenience.

Extensive beta testing was conducted by research staff at QIMR Berghofer and external consultants to ensure that there were no inconsistencies in the questionnaire and that the appropriate question skips were in place.

### Depression cases

Participants were asked “Have you ever been diagnosed with any of the following” and were presented with a list of mental health disorders with “Depression” as the first response option. We also evaluated whether participants met the DSM-5 criteria for major depressive disorder using the CIDI. The screening questions for depression were focused on the worst period of depression that a participant had experienced. Age at worst episode as well as the age at which the participant had first had a 2 week period of dysphoria or anhedonia as well as age at most recent episode were assessed. Participants were also asked to report the number of periods of at least 2 weeks of dysphoria or anhedonia they had ever had.

### Antidepressants

To assess whether participants had taken antidepressants to treat depression, the question “Have you ever taken any of the following antidepressants (even if it wasn’t for depression or anxiety)?” and were presented with a list of the 20 most commonly used antidepressants in Australia in addition to their common trade names. If they had taken one or more of the 10 most frequently prescribed antidepressants in Australia according to PBS records (sertraline, escitalopram, venlafaxine, fluoxetine, citalopram, desvenlafaxine, duloxetine, mirtazapine, amitriptyline and paroxetine), they were then asked “Why were you prescribed [name of antidepressant]”.

#### Benefits and Side-Effects of 10 most common antidepressants

Perceived effectiveness of each antidepressant medication was assessed by asking participants “How well does/did [name of antidepressant] work for you?”, with response options of “very well”, “moderately well”, “not at all well” and “don’t know”. Participants were also asked to select from a list of all side-effects that they experienced from taking each antidepressant. The list of side effects was generated from the “very common” (frequency ≥ 10%) and “common” (frequency ≥ 1% and <10%) side effects listed in the Consumer Medication Information for each antidepressant. A total of 24 side-effects were included with an “other” option also provided. Participants were also asked if they stopped taking any of the antidepressants because of side effects.

## RESULTS

### Demographics

As of 3 September 2018, questionnaire responses had been received from 20,689 participants, 75% of whom were female. The age distribution of participants, by sex, is shown for this recruitment wave in Figure 3. By the same date, saliva samples were returned by 15,807 participants (76% of the participant group). The average age of participants was 43 years ± 15 years (range 18 – 90 years), with the demographic characteristics of the cohort, as a function of recruitment method, being outlined in Table 1.

**Figure 3.**
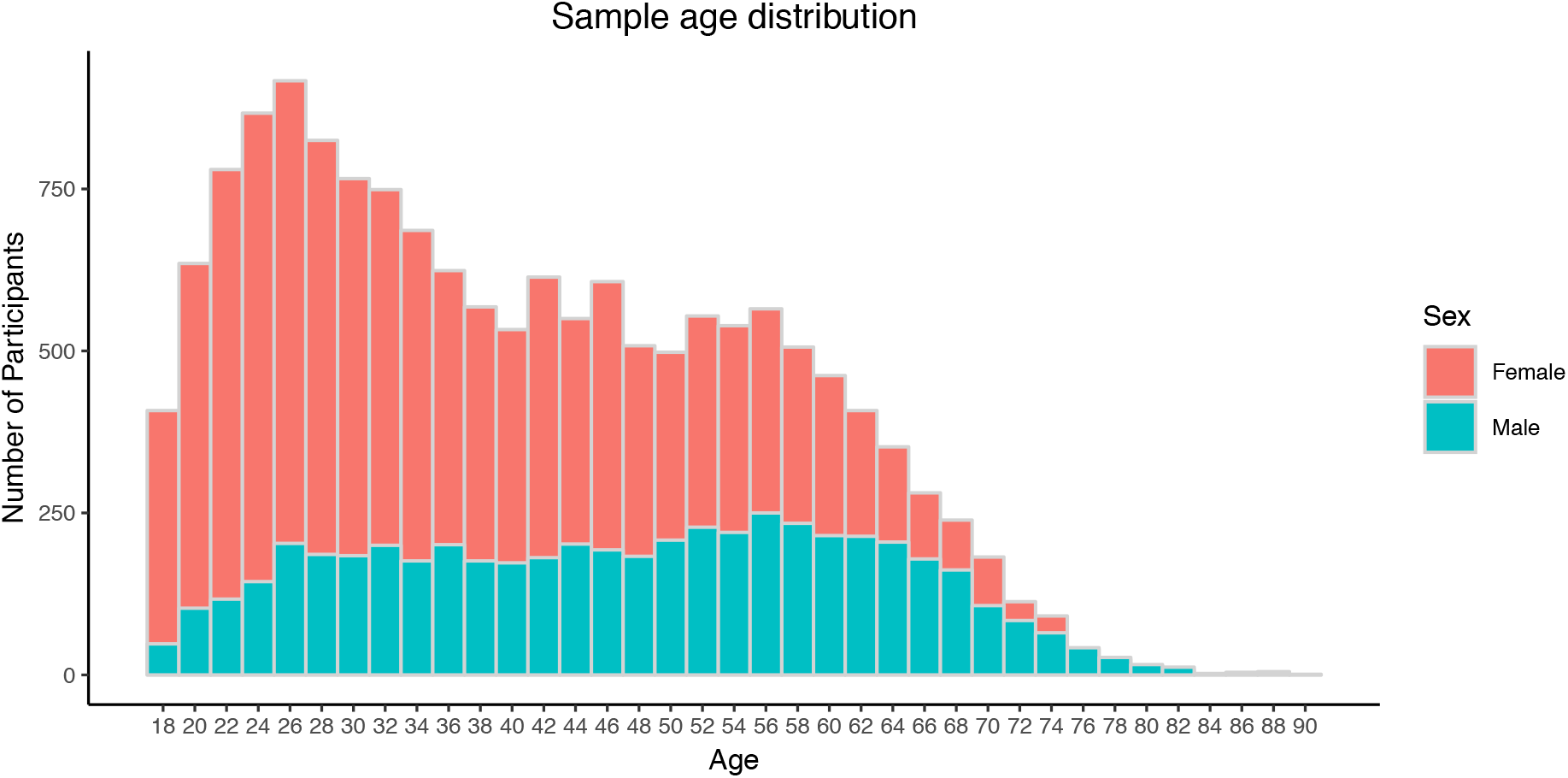
Age distribution by sex of participants in AGDS

**Table 1.**
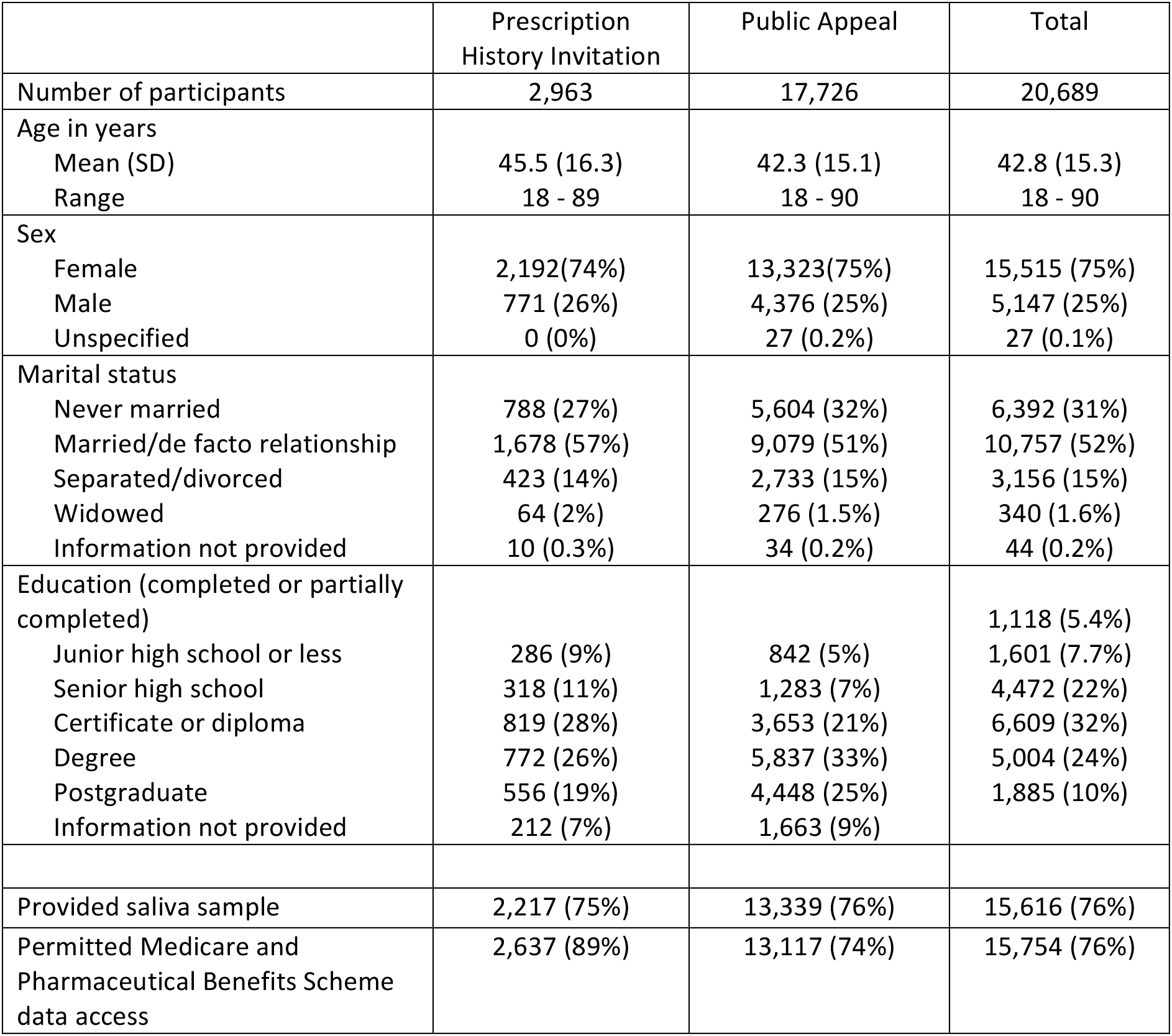
Demographic and study participation characteristics of study sample

### Mental Health Disorders

Among respondents, 98.5% reported having discussed mental health problems with a professional and 19,803 (93.4%) respondents reported having recieved a diagnosis of depression. The next most commonly reported diagnoses were Anxiety Disorder (55.0%), Posttraumatic Stress Disorder (14.0%) and Social Anxiety Disorder (11.4%). The frequency of all self-reported diagnoses is shown in Table 2.

**Table 2.** Self-reported mental health diagnostic history of study sample. Participants may report more than one diagnosis.

| Disorder | Count | Percentage of sample endorsing |
| --- | --- | --- |
| **Depression** | 19603 | 94.7 |
| **Anxiety Disorder** | 11375 | 55.0 |
| **PTSD** | 2900 | 14.0 |
| **Social Anxiety Disorder** | 2359 | 11.4 |
| **Panic Disorder** | 1960 | 9.5 |
| **Bipolar** | 1943 | 9.4 |
| **Personality Disorder** | 1200 | 5.9 |
| **Obsessive Compulsive Disorder** | 1175 | 5.8 |
| **ADD/ADHD** | 847 | 4.1 |
| **Substance Use Disorder** | 764 | 3.7 |
| **Anorexia Nervosa** | 731 | 3.6 |
| **Specific Phobia** | 724 | 3.6 |
| **Bulimia Nervosa** | 638 | 3.1 |
| **Seasonal Affective Disorder** | 582 | 2.8 |
| **Agoraphobia** | 448 | 2.2 |
| **Autism** | 331 | 1.6 |
| **Schizophrenia** | 184 | 0.9 |
| **Hoarding Disorder** | 100 | 0.5 |
| **Tourette's** | 27 | 0.1 |

### Depression diagnosed by CIDI

The DSM-5 outlines the following criteria for a depressive episode: dysphoria and/or anhedonia most of the day, nearly every day for at least 2 weeks and experiencing at least 5 out of 9 symptoms (including dysphoria or anhedonia). Consistent with the high rates of self-report diagnosis in the sample, 17,698 out of 20,165 individuals who completed the depression screening section met the criteria for a depressive episode. Additionally, 358 individuals reported not having had a 2-week period of dysphoria or anhedonia; another 1,239 reported that their symptoms persisted for less than half the day and 161 did not endorse at least 5 of the 9 symptoms required.

Mean age at onset was 22. The distribution of age at onset by sex is shown in Figure 4. The peaks between ages 10-15 and 16-20 highlight that adolescence is a peak time for developing depression. The proportion of men in each category increases with increasing age, highlighting that men are more at risk to develop depression later in life.

**Figure 4.**
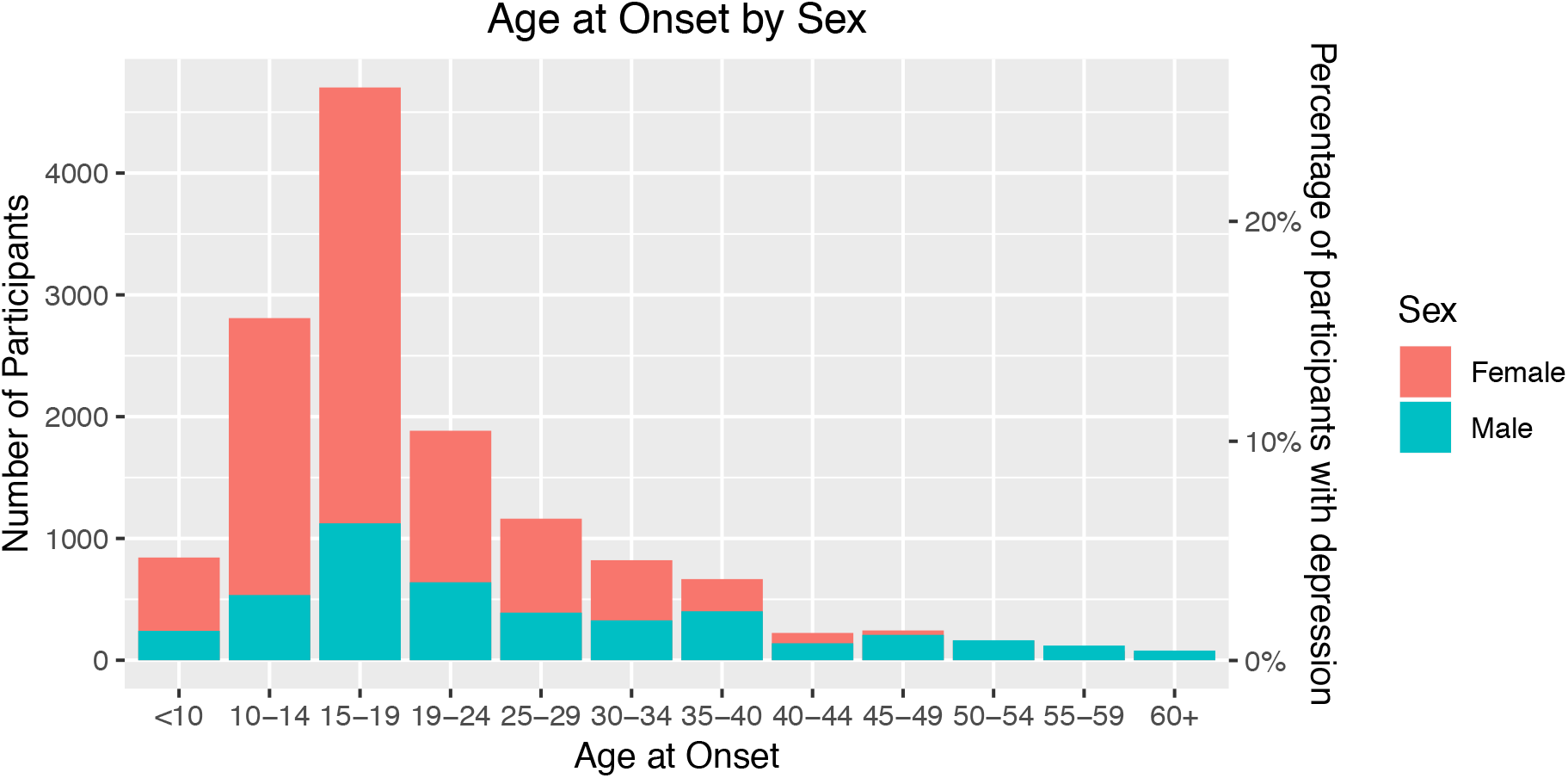
Age at onset of depression by gender

The median number of episodes reported was 6, with the most commonly reported number of periods of at least 2 weeks with depression being 13+. Only 4% of the sample report experiencing only one depressive episode (Figure 5), indicating that the sample is enriched for severe, recurrent depression.

**Figure 5.**
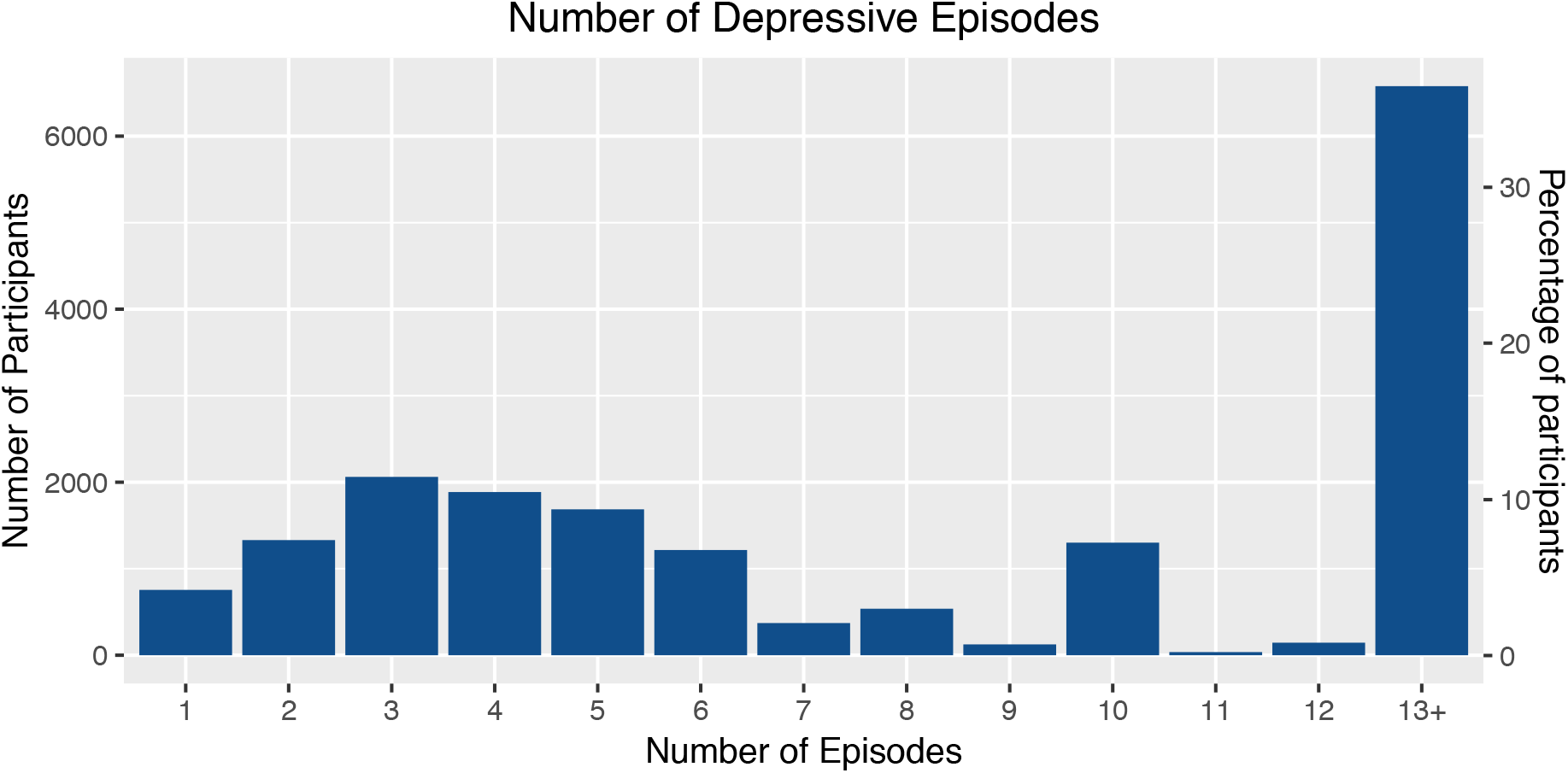
Number of reported depressive episodes among those meeting criteria for Major Depressive Disorder

The median duration of the worst episode was 12 weeks. More than 10% of the sample reported that the worst episode that they experienced was longer than a year in duration (Figure 6).

**Figure 6.**
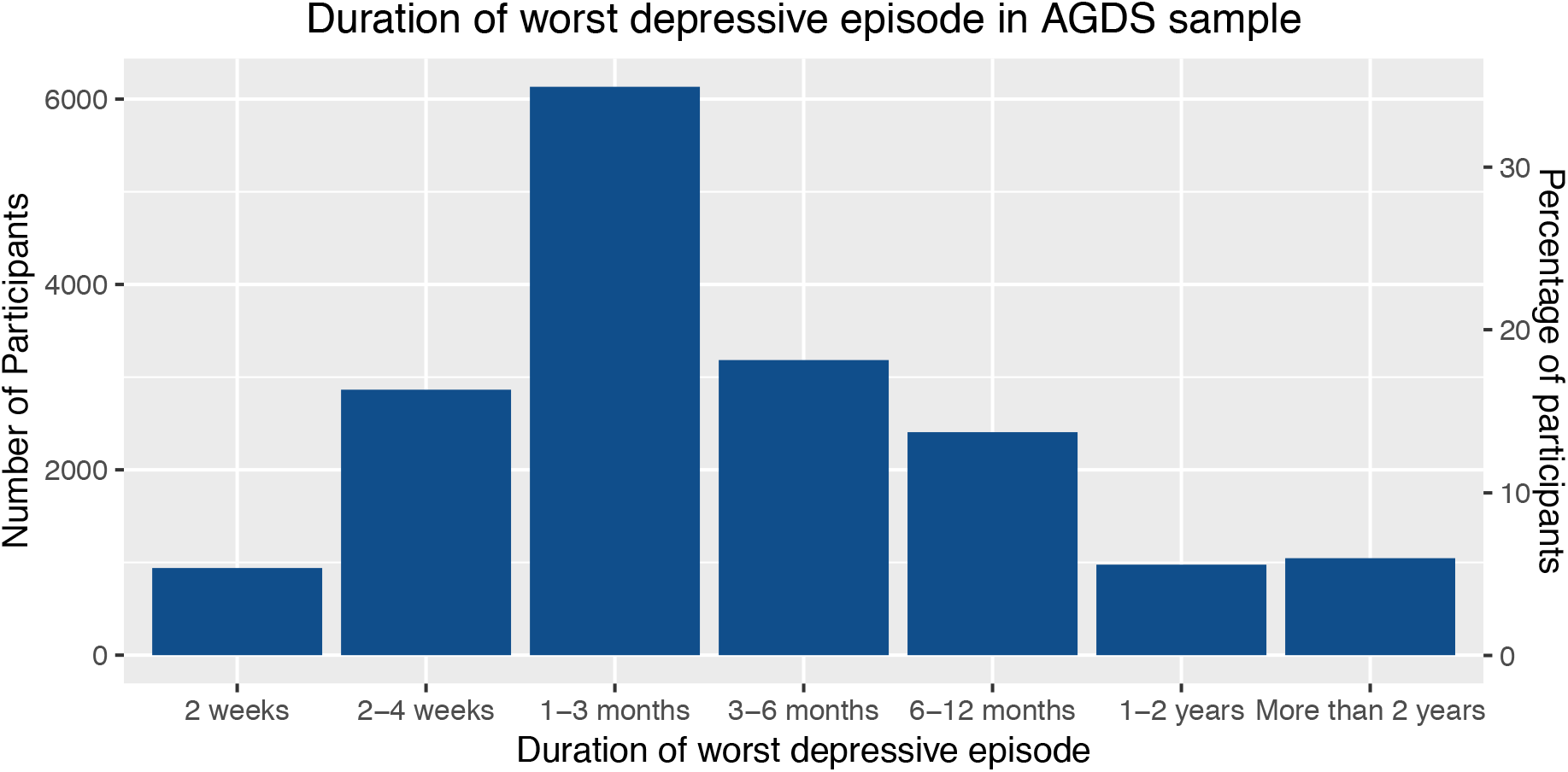
Duration of worst episode

### Family History

Out of 19,400 individuals who responded to the question about family history, 13,505 (70%) reported that a first-degree relative (parent, sibling or child) had been diagnosed with a mental health disorder. The most commonly reported diagnosis in relatives was depression, (with 11,929 individuals), followed by generalised anxiety disorder (GAD) and bipolar disorder (Figure 7).

**Figure 7.**
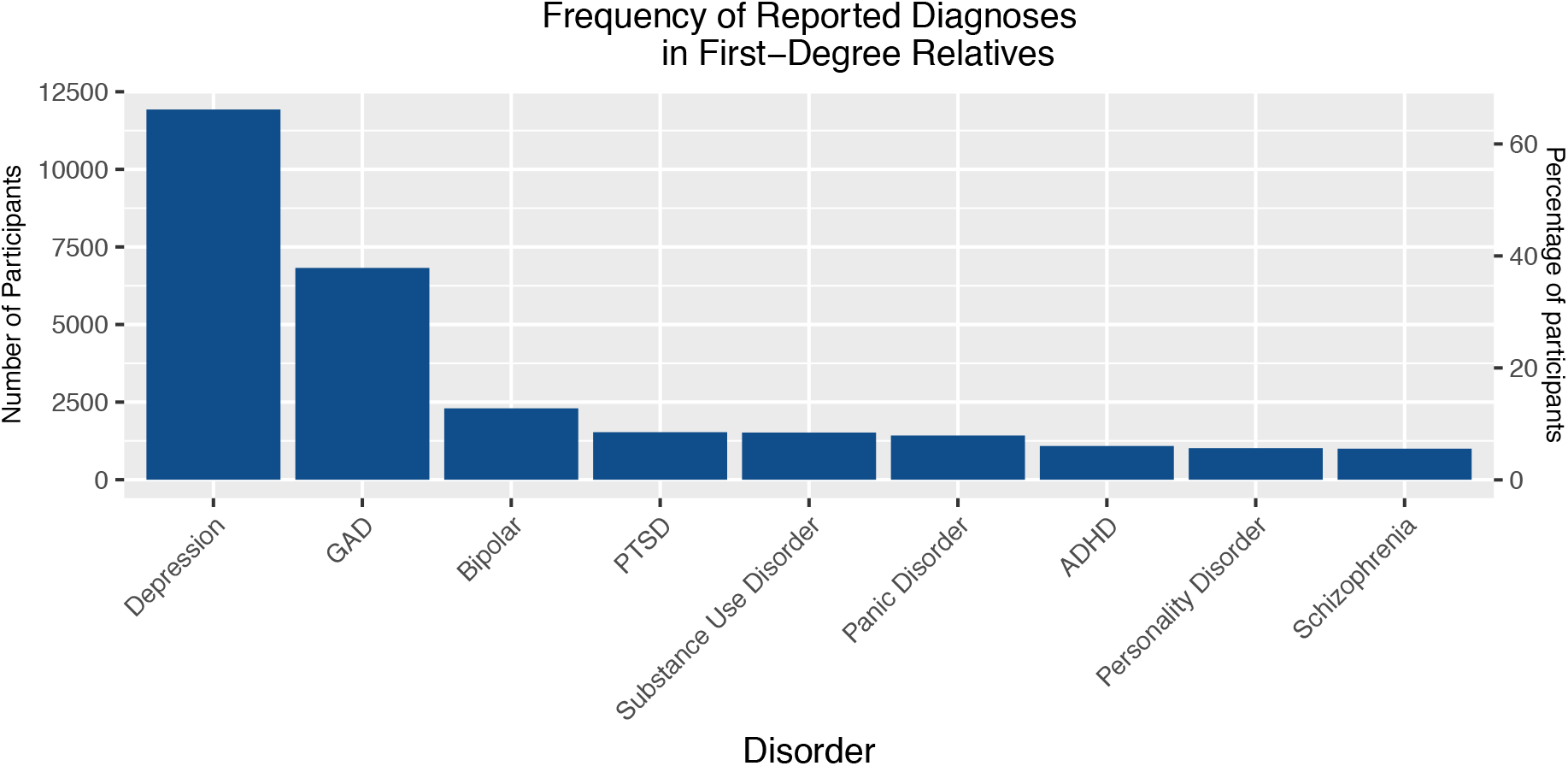
Frequency of reported diagnoses in first-degree relatives of participants

### Antidepressant Usage

A total of 95% of the sample (n = 19,585) reported taking an antidepressant. Of those reporting antidepressant use, 93% (n = 18,174) reported taking the antidepressant for depression and 51% reported taking for anxiety.

Among those taking antidepressants, the mean number of antidepressants taken was 2.75 (S.D. = 2.05, range = 1-14). Only 33% of the sample had ever taken only one antidepressant, with 42% reporting having taken 3 or more different antidepressants (Figure 8).

**Figure 8.**
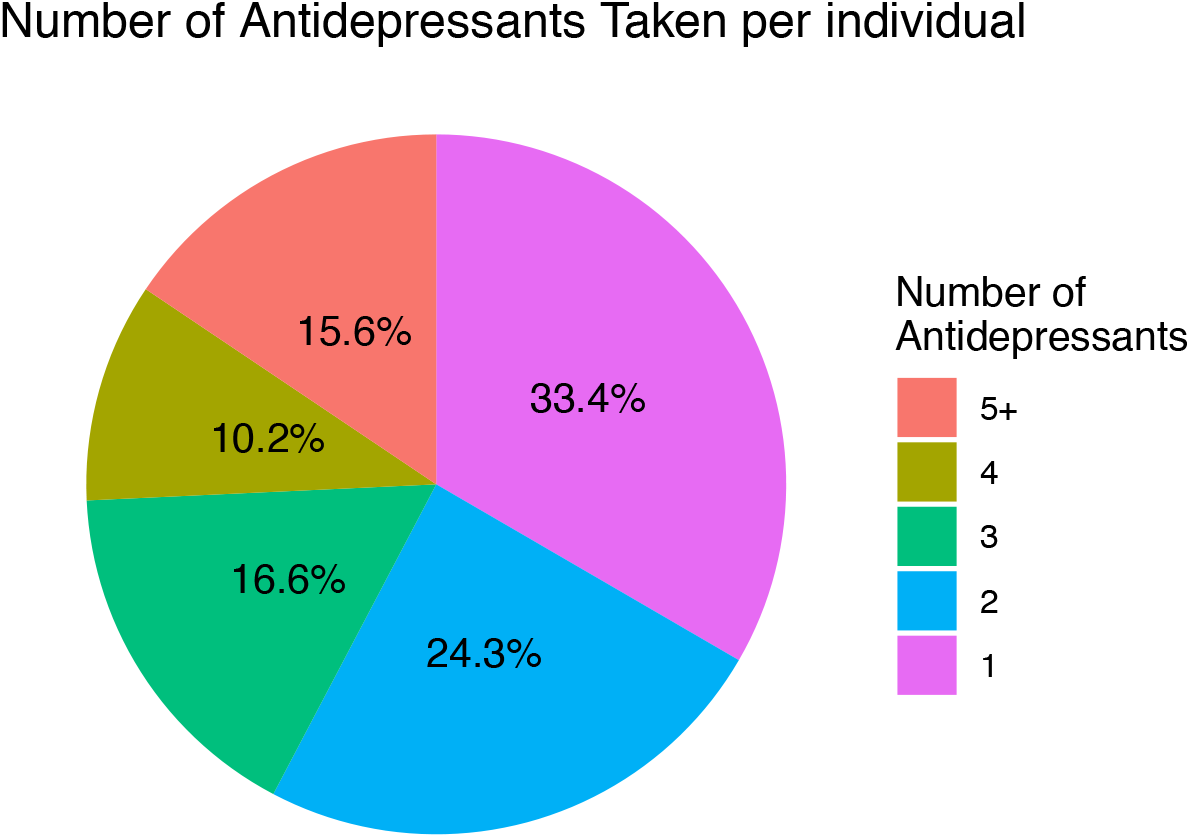
Distribution of the number of prescribed antidepressants taken by participants

**Figure 9.**
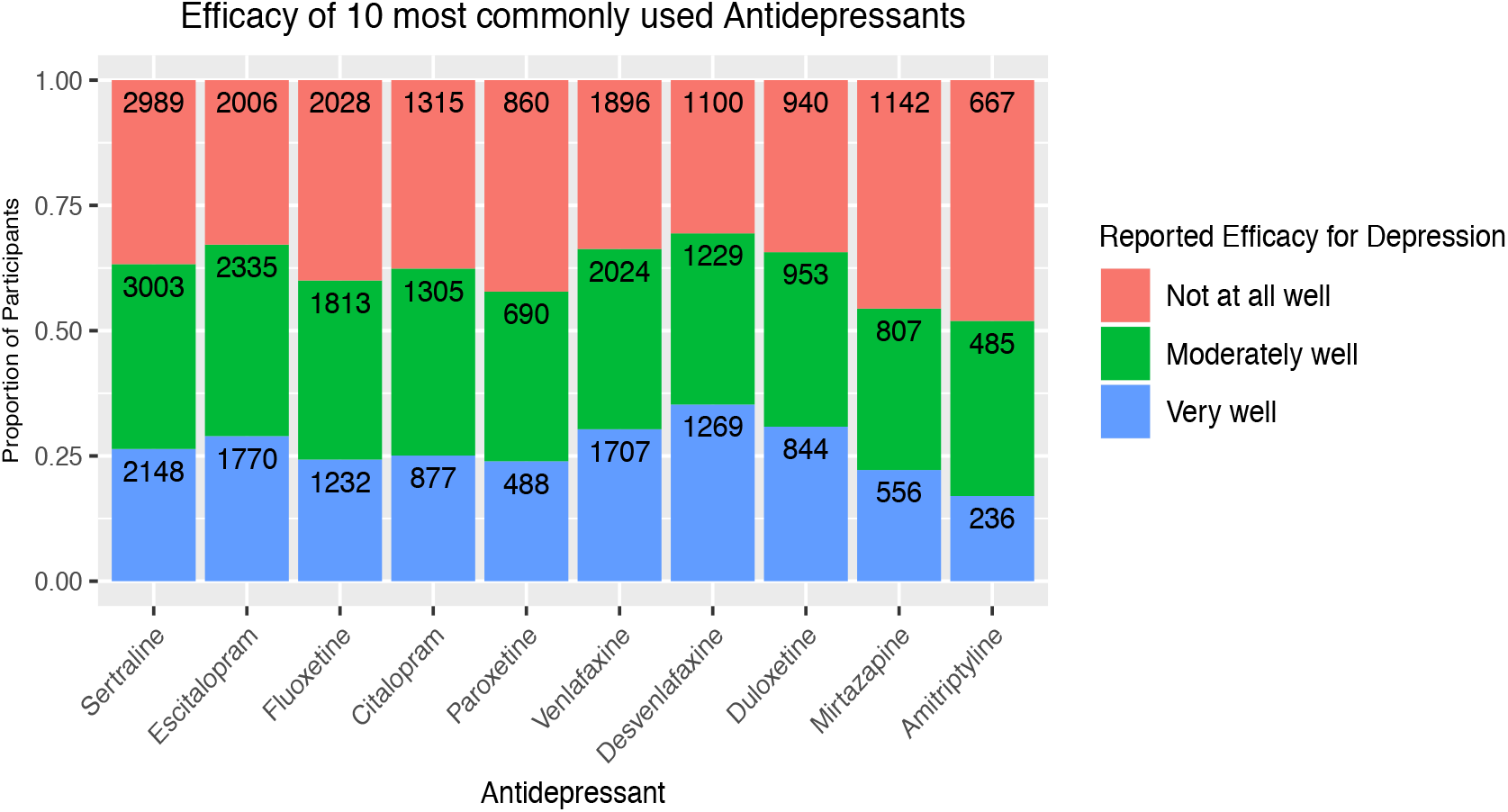
Reported efficacy of the most commonly prescribed antidepressants (numbers with each response are shown inside the bar)

For the 10 most common antidepressants listed, the number and percentage of participants with experiences of each medication are shown in Table 3. Reported effectiveness of the 10 most common antidepressants is shown in Figure 10. The rates of endorsement of the most common side-effects across the 10 most common antidepressants are shown in Table 4. More detailed analyses on the therapeutic benefits and side-effects of different antidepressants will follow in subsequent papers.

**Table 3.** Frequency of antidepressant taken in AGDS. Participants may report taking more than one antidepressant

| Antidepressant | Count | Percentage of sample endorsing |
| --- | --- | --- |
| **Sertraline** | 9132 | 44.12 |
| **Escitalopram** | 7076 | 34.19 |
| **Venlafaxine** | 6287 | 30.38 |
| **Fluoxetine** | 5823 | 28.14 |
| **Citalopram** | 4060 | 19.62 |
| **Desvenlafaxine** | 4042 | 19.53 |
| **Duloxetine** | 3168 | 15.31 |
| **Mirtazapine** | 3134 | 15.14 |
| **Amitriptyline** | 2593 | 12.53 |
| **Paroxetine** | 2471 | 11.94 |
| **Other** | 2212 | 10.69 |
| **Fluvoxamine** | 793 | 3.83 |
| **Moclobemide** | 491 | 2.37 |
| **Dothiepin** | 448 | 2.16 |
| **Nortriptyline** | 345 | 1.67 |
| **Reboxetine** | 341 | 1.65 |
| **Imipramine** | 322 | 1.56 |
| **Doxepin** | 287 | 1.39 |
| **Clomipramine** | 228 | 1.1 |
| **Tranylcypromine** | 212 | 1.02 |
| **Phenelzine** | 146 | 0.71 |
| **Mianserin** | 86 | 0.42 |
| **Never taken**<br>**antidepressants** | 976 | 4.72 |

**Table 4.** Proportion of all individuals who endorse the most common side-effects of antidepressants.

| Side Effect | Percentage of sample endorsing |
| --- | --- |
| Reduced sex drive | 35.0 |
| Weight gain | 26.3 |
| Dry mouth | 21.6 |
| Nausea | 17.6 |
| Drowsiness | 16.1 |
| Insomnia | 16.0 |
| Dizziness | 15.6 |
| Fatigue | 14.4 |
| Sweating | 14.0 |
| Headache | 14.0 |
| Suicidal thoughts | 12.3 |
| Anxiety | 11.6 |
| Agitation | 11.4 |
| Shaking | 9.3 |
| Constipation | 6.6 |
| Diarrhoea | 4.7 |
| Suicide attempt | 4.3 |
| Blurred vision | 3.9 |
| Muscle pain | 3.4 |
| Vomiting | 2.7 |
| Weight loss | 2.4 |
| Runny nose | 1.3 |
| Rash | 1.0 |

## Discussion

The Australian Genetics of Depression Study was established to recruit a large sample of participants in Australia who have experienced depression in order to better understand risk factors for depression, treatment response and side-effects. Through two modes of recruitment – government medical and pharmaceutical records and a large media campaign – more than 20,000 individuals were recruited to participate over a 2 year period. With extensive follow-up through email and, at the stage of getting saliva samples returned, phone follow-up by experienced interviewers, 76% of those enrolled returned a sample. This is one of the largest cohorts in the world with detailed information on history of depression and is a testament to the willingness of Australians to participate in medical research.

Nearly all of the study participants reported having been diagnosed or treated for depression. Using the CIDI structured interview to assess history of depression, we found that the majority of those who reported being treated for depression also meet the criteria for a depressive episode using DSM-5 criteria.

The mean age among those recruited through the media was lower than through the PBS scheme and had higher rates of university completion. This suggests that the former may be closer to a random sample from the population. It is of course unlikely that the recruitment efforts described above will generate representative samples of patients or controls, given that they rely on volunteering by as few as 5% of those asked. For GWAS it is important that cases and controls be matched for ethnicity, and this can be checked from genotyping.

These results highlight the high rate of comorbidities with depressive disorders in real-world settings (Plana-Ripoll et al., 2019). More than 60% of the sample reported having an anxiety disorder and nearly 10% reported having been diagnosed or treated for bipolar. Understanding the pattern of comorbidities and how it relates to response to treatment, emergence of side-effects (e.g greater anxiety or agitation in those with comorbid anxiety disorders), and underlying genetic variations that this scale of study can address. Specifically it will be of interest to see if there are different genetic or environmental risk factors to onset, course of illness, response to treatment or emergence of specific side-effects for those with depression and comorbid anxiety compared to depression without anxiety. In addition, we will test specific proposed subtypes of depression (e.g perinatal depression, atypical depression, chronic depression, early-onset vs late-onset depression or depression with hypomanic or brief manic features) that show evidence of distinct genetic risk factors for onset or treatment response.

Participants reported high rates of mental disorders in their first-degree relatives, highlighting the well-established genetic covariance between psychiatric disorders (Cross-Disorder Group of the Psychiatric Genomics et al., 2013). High rates of familial disorders may reflect that participants were more likely to participate in a genetic study if they have a family history or that participants shared details of the study with family members. Familial relationships will be controlled for in future genetic analyses.

Nearly half of participants reported taking 3 or more antidepressants to treat depression and thus having considerable time to improvement in symptoms. Moreover, side-effects are common and in many cases cause individuals to stop taking a drug. There is therefore an urgent need to identify risk factors for non-response to certain drugs and to reduce side effects. Not only will such advances improve the lives of patients but will also assist to reduce costs attributable to delays in achieving illness remission. Future papers will conduct finer grained analyses of response to specific antidepressants and their profile of side effects. In collecting a wide range of environmental, social and genetic data, AGDS will make a significant contribution to our understanding of variability in response and side effects.

## Acknowledgments

We are indebted to all of the participants for giving their time to contribute to this study. We wish to thank all the people who helped in the conception, implementation, beta testing, media campaign and data cleaning. We would specifically like to acknowledge Dale Nyholt for advice on using the PBS for research; Ken Kendler, Patrick Sullivan, Andrew McIntosh and Cathryn Lewis for input on the questionnaire; Lorelle Nunn, Mary Ferguson, Lucy Winkler and Natalie Garden for data and sample collection; Natalia Zmicerevska, Alissa Nichles and Candace Brennan for participant recruitment support; .Jonathan Davies, Luke Lowrey and Valeriano Antonini for support with IT aspects; Vera Morgan and Ken Kirkby for help with the media campaign. We would like to thank VIVA! Communications for their effort in promoting the study. The AGDS was primarily funded by National Health and Medical Research Council (NHMRC) of Australia grant 1086683. This work was further supported by NHMRC grants 1145645, 1078901 and 1087889.

## References

Biernacka JM, Sangkuhl K, Jenkins G, et al. (2016) The International SSRI Pharmacogenomics Consortium (ISPC): a genome-wide association study of antidepressant treatment response. Transl Psychiatry 6: e937.

consortium C. (2015) Sparse whole-genome sequencing identifies two loci for major depressive disorder. Nature 523: 588–591.

Cross-Disorder Group of the Psychiatric Genomics C, Lee SH, Ripke S, et al. (2013) Genetic relationship between five psychiatric disorders estimated from genome-wide SNPs. Nat Genet 45:984–994.

Ferrari AJ, Charlson FJ, Norman RE, et al. (2013) Burden of depressive disorders by country, sex, age, and year: findings from the global burden of disease study 2010. PLoS Med 10: e1001547.

Hodgson K, Uher R, Crawford AA, et al. (2014) Genetic predictors of antidepressant side effects: a grouped candidate gene approach in the Genome-Based Therapeutic Drugs for Depression (GENDEP) study. J Psychopharmacol 28: 142–150.

Howard DM, Adams MJ, Clarke TK, et al. (2019) Genome-wide meta-analysis of depression identifies 102 independent variants and highlights the importance of the prefrontal brain regions. Nat Neurosci.

Howard DM, Adams MJ, Shirali M, et al. (2018) Genome-wide association study of depression phenotypes in UK Biobank identifies variants in excitatory synaptic pathways. Nat Commun 9: 1470.

Investigators G, Investigators M and Investigators SD. (2013) Common genetic variation and antidepressant efficacy in major depressive disorder: a meta-analysis of three genome-wide pharmacogenetic studies. Am J Psychiatry 170: 207–217.

Li QS, Tian C, Seabrook GR, et al. (2016) Analysis of 23andMe antidepressant efficacy survey data: implication of circadian rhythm and neuroplasticity in bupropion response. Transl Psychiatry 6: e889.

Malhi GS and Mann JJ. (2018) Depression. Lancet 392: 2299–2312.

Olsen CM, Green AC, Neale RE, et al. (2012) Cohort profile: the QSkin Sun and Health Study. Int J Epidemiol 41: 929–929i.

Plana-Ripoll O, Pedersen CB, Holtz Y, et al. (2019) Exploring Comorbidity Within Mental Disorders Among a Danish National Population. JAMA Psychiatry.

Sullivan PF, Neale MC and Kendler KS. (2000) Genetic epidemiology of major depression: review and meta-analysis. Am J Psychiatry 157: 1552–1562.

Tansey KE, Guipponi M, Hu X, et al. (2013) Contribution of common genetic variants to antidepressant response. Biol Psychiatry 73: 679–682.

Tansey KE, Guipponi M, Perroud N, et al. (2012) Genetic predictors of response to serotonergic and noradrenergic antidepressants in major depressive disorder: a genome-wide analysis of individual-level data and a meta-analysis. PLoS Med 9: e1001326.

Uher R, Perroud N, Ng MY, et al. (2010) Genome-wide pharmacogenetics of antidepressant response in the GENDEP project. Am J Psychiatry 167: 555–564.

Whiteford HA, Degenhardt L, Rehm J, et al. (2013) Global burden of disease attributable to mental and substance use disorders: findings from the Global Burden of Disease Study 2010. Lancet 382: 1575–1586.

Wray NR, Pergadia ML, Blackwood DH, et al. (2012) Genome-wide association study of major depressive disorder: new results, meta-analysis, and lessons learned. Mol Psychiatry 17: 36–48.

Wray NR, Ripke S, Mattheisen M, et al. (2018) Genome-wide association analyses identify 44 risk variants and refine the genetic architecture of major depression. Nat Genet 50: 668–681.

